# Global determinants of mRNA degradation rates in *Saccharomyces cerevisiae*

**DOI:** 10.1101/014845

**Authors:** Benjamin Neymotin, Victoria Ettorre, David Gresham

**Affiliations:** Center for Genomics and Systems Biology, Department of Biology, New York University

## Abstract

Degradation of mRNA contributes to variation in transcript abundance. Studies of individual mRNAs show that *cis* and *trans* factors control mRNA degradation rates. However, transcriptome-wide studies have failed to identify global relationships between transcript properties and mRNA degradation. We investigated the contribution of *cis* and *trans* factors to transcriptome-wide degradation rate variation in the budding yeast, *Saccharomyces cerevisiae*, using multiple regression analysis. We find that multiple transcript properties are associated with mRNA degradation rates and that a model incorporating these factors explains ∼50% of the genome-wide variance. Predictors of mRNA degradation rates include transcript length, abundance, ribosome density, codon adaptation index (CAI) and GC content of the third position in codons. To validate these factors we studied individual transcripts expressed from identical promoters. We find that decreasing ribosome density by mutating the translational start site of the *GAP1* transcript increases its degradation rate. Using variants of GFP that differ at synonymous sites, we show that increased GC content of the third position of codons results in decreased mRNA degradation rate. Thus, in steady-state conditions, a large fraction of genome-wide variation in mRNA degradation rates is determined by inherent properties of transcripts related to protein translation rather than specific regulatory mechanisms.

## INTRODUCTION

Alterations in the abundance of mRNA result from changes in both the rate of transcript synthesis and the rate of transcript degradation. Synthesis and degradation of mRNAs is critical for control of gene expression and cell survival as ablation of either process results in rapid loss of viability (1, 2). The *cis* and *trans* factors that control rates of mRNA synthesis have been extensively studied in many systems (reviewed in (3)). By comparison, far less is known about factors that control rates of mRNA degradation. A complete understanding of gene expression regulation requires identification of the sources of variation in mRNA degradation.

Our understanding of the mechanisms by which mRNAs are degraded (reviewed in (4)) is largely the result of studies of specific transcripts (5–7). These studies have shown that mRNA degradation is controlled by *cis* factors, including sequence elements in the coding (8, 9) and untranslated (10, 11) regions, as well as factors in *trans*, including RNA binding proteins (12, 13) and non-coding RNAs (14). However, the extent to which these different factors impact global patterns of mRNA degradation remains unclear.

Genome-wide mRNA degradation rates have been determined for a number of organisms including bacteria (15), plants (16), flies (17), mouse (18) and human cell lines (19). In the budding yeast, *Saccharomyces cerevisiae*, genome-wide mRNA degradation rates have been measured using a variety of methods including transcriptional inhibition (20–22), genomic-run-on (23), and metabolic labeling (24, 25). In general, the concordance between different global studies of mRNA degradation rates is poor likely due to a combination of technical and biological sources of variation. Recently, we introduced <u>R</u>NA <u>A</u>pproach <u>t</u>o <u>E</u>quilibrium <u>Seq</u>uencing (RATE-seq), which combines 4-thiouracil (4-tU) labeling and RNA-seq for determination of genome-wide *in-vivo* mRNA degradation rates (26). Using approach to equilibrium labeling kinetics and non-linear regression, RATE-seq overcomes several problems with existing methods providing improved accuracy of mRNA degradation rates estimates, and a measure of the precision with which the rates are measured, in steady-state conditions.

Despite discrepancies in estimates of mRNA degradation rates among different studies, three consistent features have been demonstrated across multiple transcriptome-wide datasets. First, there is variation in the rates at which different transcripts are degraded: some vary by as much as an order of magnitude. Second, transcripts for genes encoding functionally-related products have similar degradation rates (20, 26, 27). Third, no single property of transcripts explains the observed variation (20, 25, 24). This latter point suggests that either the causative factors are obscured in genome-wide studies, or a combination of different factors affect rates of degradation that have transcript specific effects. Potential properties of transcripts that might impact their rate of degradation include transcript length, GC content, transcript abundance, codon usage and folding properties. However, testing the effect of any single property of transcripts on global degradation rates is inherently challenging as each parameter can vary independently across transcripts. At the same time many transcript properties are correlated with each other making it difficult to identify causative factors. Thus, to identify determinants of variation in mRNA degradation rates all known properties of transcripts must be considered simultaneously and experimental designs that modulate a single property are required for validation experiments.

Here, we analyzed mRNA degradation rates in *S. cerevisiae* using multiple regression analysis (28, 19) and genome-wide datasets that have defined gene specific parameters. These include protein levels (29), protein half-life (30), RNA abundance (31), transcription rates (32), UTR lengths (33), ribosome density (34), association with RNA binding proteins (35) and the function of the encoded product. A multiple regression model applied to mRNA degradation rates determined using RATE-seq accounts for 50% of the variation in mRNA degradation rates. Although less variation is explained for multiple regression models applied to other genome-wide mRNA degradation datasets, many predictors are significant in multiple datasets, suggesting that they are reproducible transcript properties that impact degradation rates. These features include ribosome density, codon adaptation index, and GC content of the wobble position in a codon (GC3) suggesting that protein translation is closely related to mRNA degradation.

Using experimental studies of individual transcripts, we show that changing ribosome density affects both the mRNA degradation rate and steady state levels. Using GFP coding sequence variants that differ only in their GC3 content we show that coding sequence affects mRNA degradation. Increasing the GC3 content increases the stability of mRNAs resulting in increased steady-state level. Our results suggest that mRNA degradation is determined by multiple factors, many of which are intimately linked to protein translation.

## MATERIALS AND METHODS

### Plasmid construction

Plasmid pCM188 (36) was used as the backbone for all plasmids. This CEN4 plasmid contains the *URA3* gene, a constitutively expressed tetracycline transactivator, and a multiple cloning site with a *CYC1* TATA region upstream, all under control of two copies of the tetracycline operator. Transcription of the gene is repressed in the presence of tetracycline or its derivatives, including doxycycline. Plasmids DGP147, DGP148, DGP149, and DGP231 are pCM188 with degenerate forms of GFP (37) ranging in GC content in the third position of each codon as 0.71, 0.38, 0.60, and 0.67 respectively (GFP4, GFP1, GFP2, GFP3). The coding sequence of each GFP was cloned into the BamHI and NotI sites. Plasmid DGP217 is pCM188 with the *GAP1* gene + 3’UTR cloned into the BamH I and Not I sites. Plasmid DGP218 is the same as DGP217, except the start codon of *GAP1* has been mutated to GTG.

### Strains and Growth Conditions

The laboratory strain, FY3 (*MATa ura3-52*) which is isogenic to S288C was used for all experiments. DGY696, 697, 698, and 1281 have plasmids DGP147, 148, 149, and 231 respectively. DGY1193 and 1194 carry plasmids DGP217 and DGP218 respectively, with the *GAP1* locus deleted from start to stop with the KANMX4 cassette. For all experiments, a single colony for each strain was inoculated in synthetic complete media without uracil, to maintain selection of the plasmids. Studies of the *GAP1* transcript were performed in nitrogen limiting media with proline as the limiting nitrogen source, as described in (38). Saturated cultures from overnight cultures were back-diluted 1:50 into media of the same composition. Cells were allowed to grow for 5.5 hrs (∼2.5 doublings) before transcription was inhibited with doxycycline at a final concentration of 10μg/mL. Cells were collected by filtration on nitrocellulose membranes and snap frozen in liquid nitrogen.

### RNA processing and qRT-PCR analysis

RNA was extracted using the hot phenol-chloroform method as in (26). Purified RNA was then treated with RQ1 DNAse according to manufacturer recommendations. Reverse transcription was performed using random hexamers and MMLVRT enzyme. Quantitative Reverse Transcription PCR (qRT-PCR) was performed using the SYBR Green system and a Roche Light Cycler. RNA levels were quantified in comparison to the HTA1 housekeeping gene, which is unaffected by Doxycycline addition. Ratios were calculated using the formula: Y=2H^(-(HTA1_ct-Gene-ct))^ with ct being the calculated cycle threshold. RNA levels from each time point were normalized to t=0, which was set to 1. All analyses with error bars are the mean +/− the standard error for 3–6 biological replicates. Values without error bars are the average of two replicates. Primer sequences had amplification efficiencies of at least 95% on RT products. The amplified product for all GFP strains begins between position 463 and 586 of the transcript. The amplified products are all between 80 and 120 base pairs long. The sequences used for qRT-PCR analysis are as follows: *HTA1* (Forward: 5’- GCTGGTAATGCTGCTAGGGATA-3’, Reverse: 5’- TTACCCAATAGCTTGTTCAATT-3’), *GFP2, GFP4* (Forward: 5’-TTGCCGGATAACCACTACCT-3’, Reverse: 5’-CCTGCTGCAGTCACAAACTC-3’), *GFP3* (Forward: 5’-GCCGATAAGCAGAAGAATGG-3’, Reverse: 5’-TGTTGATAATGGTCCGCAAG-3’), GFP4 (Forward: 5’-CGACCATTACCAGCAGAACA-3’, Reverse: 5’- GGGTCCTTTGACAGAGCAGA-3’), *GAP1-ATG, GAP1-GTG* (Torward: 5’-TTTGTTCTGTCTTCGTCAC-3’, Reverse: 5’-CTCTACGGATTCACTGGCAGCA-3’)

### Multiple Regression Analysis

For multiple regression analysis we used degradation rates rather than half-lives, which are typically log-normally distributed. To minimize the effects of extreme outliers we removed values more than 1.5 times the interquartile range. Degradation rates for most datasets were calculated as ln(2)/t_half-life_, except in (25) and (26) where the effects of dilution as a result of cellular growth was also considered. Transcript counts were from (31) and estimated based on an assumption of ∼60,000 mRNA/cell (39). Protein per mRNA was calculated as the values from (29) divided by the values for counts. Codon adaptation index (40) was calculated for each transcript based on the codon frequency tables in the seqinr package in R (R core team). To normalize the data we log transformed the predictor variables (log_10_(Variable) or log_10_(Variable +1)), except for GC content of each codon position, which is approximately normal in distribution, and **Δ**G, which is negative in value. For categorical variables, transcripts were classified as bound by an RNABP based on data from (35), and as present or absent from a Gene Ontology group based on GO SLIM files downloaded from SGD.

In a linear multiple regression model (Crawley 2012), a parameter of interest is modeled as being dependent on two or more predictors. We use the degradation rate constant as the parameter of interest, and the other measurements as predictors. To build the model, we followed two separate approaches. In the first approach, we first determined the p-value of the pair wise correlation of each predictor to degradation rate. This indicates whether the regression coefficient is significantly different from zero, and whether or not the predictor has any effect on degradation rate. Next we included all predictors that have a p-value less than 0.05 into the multiple regression models. We then performed stepwise deletion of terms by removing sequentially the predictors with the highest p-value. The final model is then the reduced model where only significant terms remained. We obtained the same result using the step function in R, which reduces models based on Akaike’s Information Criterion (AIC). In a second approach, we first calculated the significance of each predictor when it is the only one in the model, as above. We then started adding to the model based on the predictors with the lowest p-value. With each additional term we checked to see that all of the terms in the model were significant. If a new term was added and it was not significant, we removed it from the model. If a new term was added and a different predictor lost its significance, we tested a model with either the new predictor or the one that lost significance, and retained the one that explained more variation. We did not add terms that were insignificant in the pair wise correlation with degradation rate. Both approaches gave similar results. Model diagnostics suggest there is no obvious curvature or patterns in terms of increase or decrease in variance as a function of fitted values (Figure S6). There is also minimal curvature in the normal Q-Q plot, suggesting the model follows linearity (Figure S6).

### R functions and packages

We performed all analyses using R (R Core Team 2013) and several open source packages. In addition to custom written functions in R, we also used functions from the following packages: *TeachingDemos, Biostrings, LSD, stringi, GeneRfold and seqinr*.

## RESULTS

### Multiple transcript properties affect global mRNA degradation rates

Previous studies have found evidence for the effect of specific properties of transcripts on the degradation rate of individual transcripts (10, 8, 11, 12). We tested the relationship between globally measured transcript features (Table 1) and genome-wide mRNA degradation rates determined using RATE-seq (Table S1). We find that several transcript features are significantly correlated with mRNA degradation rates (Figure 1A and Table S2). The most significant single feature predictive of mRNA degradation rates measured using RATE-seq is the length of the coding sequence, which explains almost 30% of the variance. The folding energy (**Δ**G) is also significantly associated with mRNA degradation rates, which may be due to the fact that folding energy and coding sequence length are highly correlated. Several features related to the translation of transcripts are also significantly associated with mRNA degradation rates including ribosome density, the codon adaptation index and the GC3 content. We also tested whether the function of the encoded product is predictive of mRNA degradation rate and found that functional assignment using gene ontology (GO) terms explains a significant fraction of the variation (Figure 1A). This is consistent with the observation that transcripts encoding proteins in similar functional categories degrade with similar kinetics (20, 26, 27). In addition, association with specific mRNA binding proteins also explains a significant fraction of variation in mRNA degradation rates (Figure 1A). These results suggest a relationship between several transcript features and mRNA degradation rates. We observed similar relationships between these predictors and mRNA degradation rates measured using other methods (Figure S1) suggesting that some of these relationships are reproducible despite the poor agreement in mRNA degradation rates among different studies.

**Figure 1.**
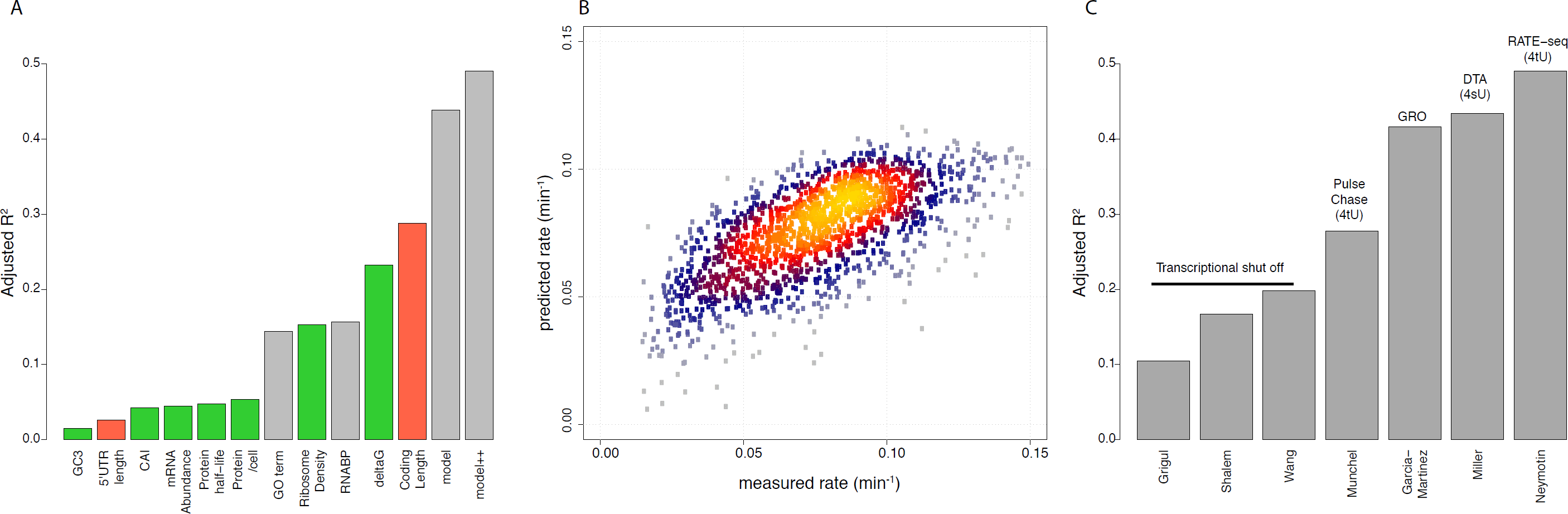
Multiple factors are associated with variation in mRNA degradation rates. (**a**) Individual predictors explain different amounts of the variation in mRNA degradation rates determined using RATE-seq. Multiple regression models including continuous variables (model) and continuous and categorical variables (model++) explain around half the variation in mRNA degradation rates. Positive correlations are indicated in red whereas negative correlations are indicates in green. Values corresponding to ANOVA and multiple regression are in blue. (**b**) A comparison of modeled mRNA degradation rates with measured mRNA degradation rates shows that the model behaves similarly across the entire range of mRNA degradation rates. (**c**) Multiple regression models applied to published mRNA degradation rates explain different amounts of the variation. Less variance can be explained for mRNA degradation rates that rely on transcriptional inhibition. Genomic Run On is indicated with GRO. Dynamic Transcriptome Analysis is indicated with DTA. Labeling studies using thiouracil or thiouridine are indicated in parentheses with 4tU and 4sU respectively.

**Table 1.** Parameters included in model.

| **Predictor** | **N** | **Units** | **Transformation** | **Reference** |
| --- | --- | --- | --- | --- |
| Coding Length | 5850 | nucleotides | $\log_{10}(\text{nucleotides})$ | SGD |
| 3'UTR length | 4950 | nucleotides | $\log_{10}(\text{nucleotides}+1)$ | SGD/ (33) |
| 5'UTR length | 4345 | nucleotides | $\log_{10}(\text{nucleotides}+1)$ | SGD/ (33) |
| 5'UTR GC content | 4345 | proportion | none | SGD/(33) |
| 3'UTR GC content | 4911 | proportion | none | SGD/(33) |
| Abundance | 5488 | rpM | $\log_{10}((\text{rpM}/\text{total reads}) * 60,000)$ | (31) |
| Protein/ cell | 3818 | protein/cell | $\log_{10}(\text{protein}/\text{cell})$ | (29) |
| Ribosome density | 5269 | rpkM | $\log_{10}(\text{rpkM}+1)$ | (34) |
| Transcription rate | 4346 | molecules/min | $\log_{10}(\text{molecules}/\text{min})$ | (32) |
| Coding GC position 1 | 5850 | proportion | none | SGD |
| Coding GC position 2 | 5850 | proportion | none | SGD |
| Coding GC position 3 | 5850 | proportion | none | SGD |
| Protein half-life | 3164 | min | $\log_{10}(\text{min})$ | (30) |
| deltaG | 5850 | kcal/mol | none | SGD/GeneRFold |
| CAI | 5850 | relative scale | $\log_{10}(\text{relative scale})$ | CAI function in R |
| Protein/ mRNA | 3583 | protein/cell/<br>transcript | $\log_{10}(\text{protein}/\text{cell}/\text{transcript}+1)$ | (29, 31) |

Although many transcript properties are correlated with each other (Figure S2), some properties show no correlation and therefore may exert independent and differential effects on the rate of mRNA degradation. Therefore, we used multiple regression analysis to model the contribution of multiple transcript features to variation in mRNA degradation rates simultaneously (**methods**). We initially built a model incorporating all factors and used sequential reduction to arrive at a minimal model (**methods**). We find that the explained variation when multiple transcript properties are included exceeds the variance explained by any single factor suggesting that degradation rates are determined by a combination of transcript features (“model” in Figure 1A). When the categorical factors of gene function and binding to specific proteins, are included 50% of the variance in mRNA degradation rates can be explained (“model++” in Figure 1A). Thus, the rates predicted by a multiple regression model are in good agreement with experimentally determined rates (Figure 1B). Models incorporating all features explain significant fractions of the variation reported in other mRNA degradation datasets, albeit with reduced explanatory power (Figure 1C). Interestingly, we find that models applied to mRNA degradation rates measured using transcriptional inhibition tend to explain much less variation than models applied to RNA degradation rates measured with different metabolic labeling methods.

Our model suggests an inverse relationship between mRNA degradation rate and translational elongation rates, as measured by Codon Adaptation Index (CAI) and ribosome density. Translation elongation rates are slowed during peptide bond formation for proline residues (41), and particularly when multiple prolines are encoded sequentially. To further investigate the role of translation elongation in mRNA degradation, we classified transcripts based on presence or absence of at least four sequential proline codons. Consistent with our multiple regression model, transcripts rich in proline degrade more rapidly than the rest of the transcriptome (Figure 2A). Interestingly, stretches of proline codons are also associated with overall lower protein expression levels (Figure 2B).

**Figure 2.**
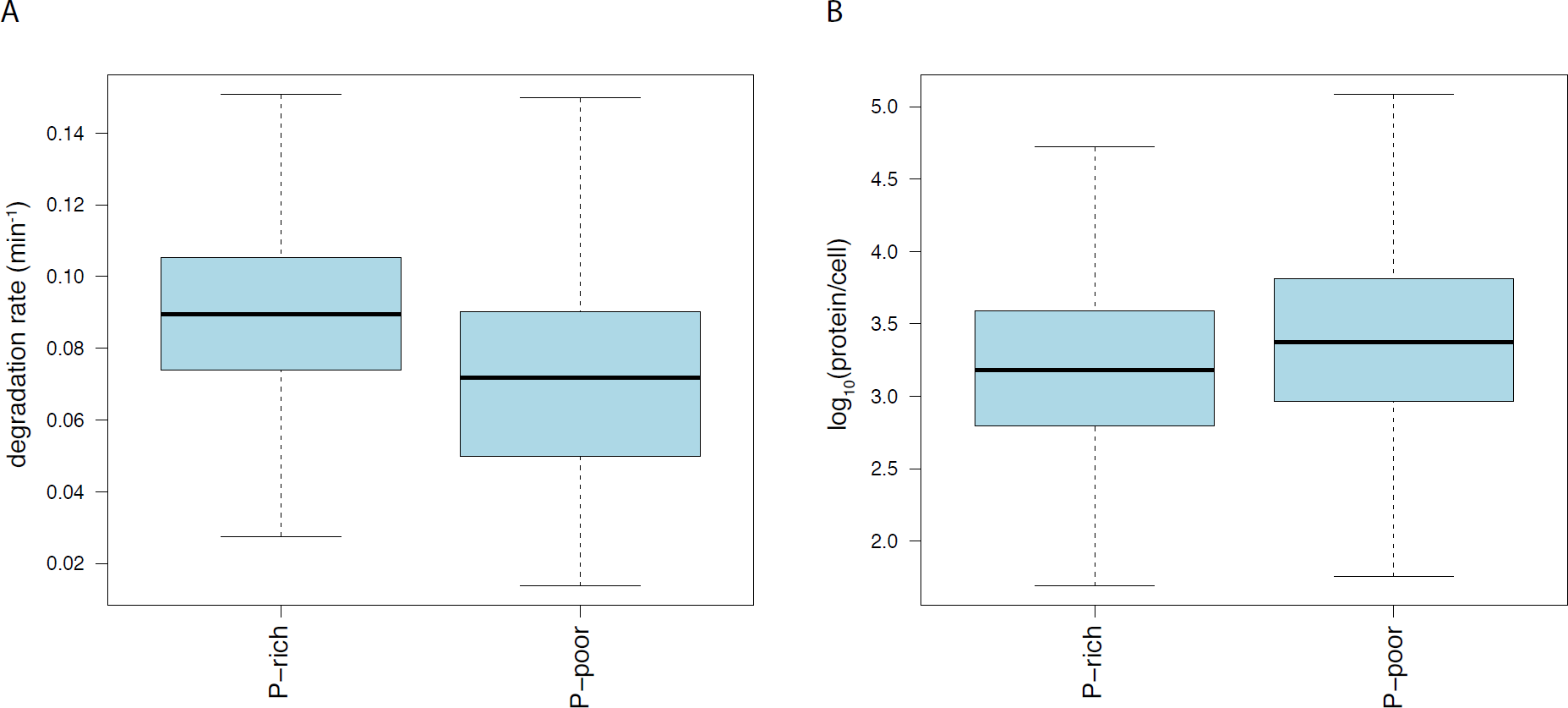
The presence of multiple proline codons affects degradation rates and protein production. (**a**) Proline-rich proteins, which are translated more slowly, tend to have more stable transcripts than proline-poor proteins. (**b**) The steady-state abundance of poly-proline containing proteins is reduced compared to the global distribution of protein abundances.

#### Ribosome density affects mRNA degradation rate

Our regression analysis suggests that different aspects of translation affect mRNA degradation rates. Ribosomes are generally thought to protect mRNAs from degradation (4). Consistent with previous analyses (42), we find that increased ribosome density is associated with decreased rates of mRNA degradation (Figure S3). To experimentally reduce the density of ribosomes on specific transcripts, we mutated the start codon of an endogenous transcript *GAP1*, which encodes the general amino acid permease, from ATG to GTG and placed it under control of doxycycline-repressible promoters (36). Addition of doxycycline has little effect on cellular physiology and no detectable effect on global gene expression (43). ATG start codons are required for the small ribosomal subunit to recruit the large ribosomal subunit for fully formed ribosomes. Mutation of ATG to GTG is expected to reduce the number of ribosomes bound to an mRNA, but not eliminate ribosome binding entirely as a downstream ATG may serve as a start codon for 80s ribosome formation.

We tested the *GAP1* transcripts for alteration in degradation kinetics as a function of start codon mutation. In the absence of a start codon, we find that the *GAP1* transcript is significantly decreased in steady state mRNA abundance (Figure 3A) and the transcript degrades more rapidly upon transcriptional inhibition (Figure 3B and Figure 3C). Our results suggest that a decrease in ribosome density increases the degradation of the *GAP1* transcript, consistent with the global trend detected in our regression model

**Figure 3.**
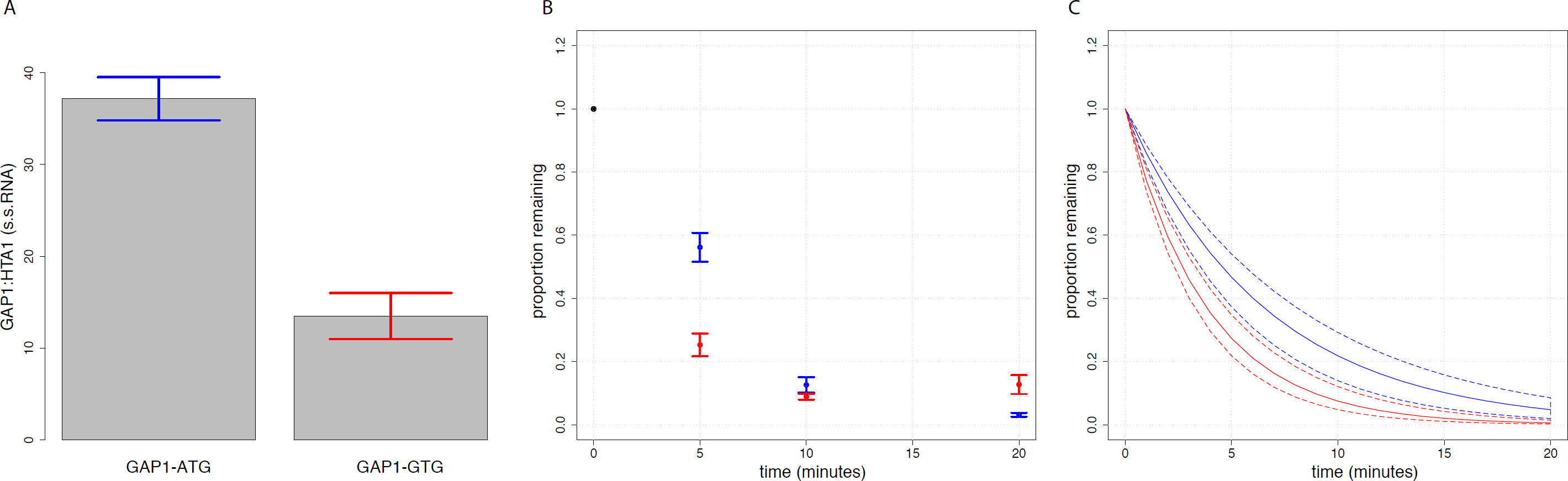
The effect of ribosome density on *GAP1* mRNA degradation rate. (**a**) Mutation of the start codon in GAP1 from ATG to GTG results in a reduced steady state transcript level (**a**) and (**b**) increased mRNA degradation. In (b) we show the average value for each time point +/− SEM. In (**c**) we show bootstrapped confidence intervals for the regression of all data points. Solid lines indicate the line of best fit and dotted lines indicate confidence intervals. In blue is the transcript with a normal start codon. In red is the transcript with a mutated start codon.

#### Decreased GC content of the third codon position increases the rate of mRNA degradation

Our multiple regression model predicts that factors involved in translation including ribosome density, codon adaptation index and the GC3 content, contribute to variation in mRNA degradation rates (Figure 1A). GC3 content affects mRNA abundance and mRNA degradation in *E. coli* (37). By studying a small set of transcripts in mammalian cells, GC3 content was also found to affect mRNA levels, but not degradation rates (44) implying that mRNA synthesis or processing must underlie differences in mRNA levels. However, a more recent genome-wide study found evidence that decreased GC3 content is correlated with increased mRNA degradation rates (19).

To study the contribution of GC3 content to variation in mRNA degradation rates we used GFP constructs that differ in sequence at synonymous sites only (37). We studied four GFP transcripts that span a range of GC3 content (Figure 4A). Changes in GC3 content also results in overall variation in total GC content (Figure 4A). Coding sequences were placed under control of the identical doxycycline-regulated promoter and engineered to have the same UTRs (Figure 4B). We confirmed that all four constructs result in functional GFP expression (data not shown).

**Figure 4.**
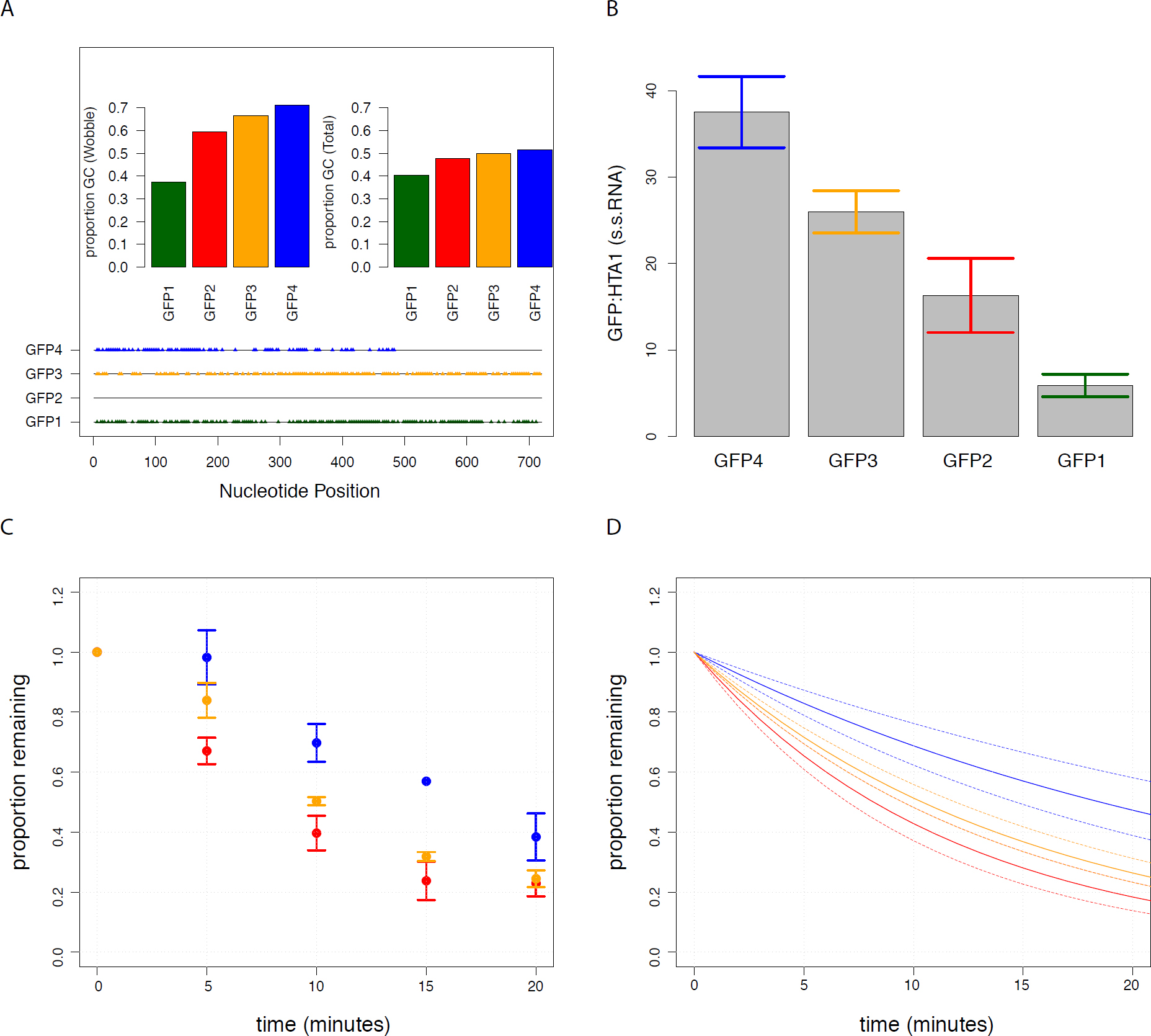
GC3 content affects degradation kinetics and steady state levels. (**a**) Four GFP transcripts containing synonymous mutations span a range of GC3 content (left) and overall GC (right). Alignment of each GFP relative to GFP2. Positions of similarity in sequence are depicted by gray line and differences are in triangles. Colors indicate different GFP constructs. (**b**) Differences in GC3 content affect steady state levels of transcripts. (**c,d**) GFP2, GFP3 and GFP4 degrade in a GC3-dependent manner. In (**c**) we show the average value for each time point +/− SEM. In (**d**) we show bootstrapped 95% confidence intervals for the regression of all data points. Solid lines indicate the line of best fit and dotted lines indicate confidence intervals.

As all coding sequences are expressed from an identical promoter differences in steady state mRNA abundance must result from differences in degradation, synthesis or processing. We find that steady-state mRNA levels vary with GC3 content with the highest GC3 content resulting in the highest steady-state mRNA abundance (Figure 4B) consistent with observations in mammalian cells (44). Following addition of doxycycline to repress transcription initiation, we confirmed that three of the four transcripts degrade differentially in a GC3-dependent manner (Figure 4C and 4D), consistent with our multiple regression prediction. Using the measured steady-state abundances and degradation rates for the three transcripts, we estimated synthesis rates. All three strains have similar estimated rates of synthesis consistent with differences in degradation rates being the primary determinant of differences in steady-state mRNA levels. We find that a fourth construct, which has much lower GC3 content, and the lowest steady-state abundance does not significantly differ in its mRNA degradation rate from the second lowest GC3 (Figure S4). This may reflect a limitation of the statistical power of our assay and the fact that other factors are likely to interact with the GC3 effect.

## DISCUSSION

The abundance of mRNAs is determined by both the rate at which they are synthesized and the rate at which they are degraded. Previous studies of individual transcripts have determined *cis* and *trans* factors that affect mRNA degradation rates (4, 45). However, variation in genome-wide mRNA degradation rates cannot be explained by any single property of transcripts. In this study we sought to construct a comprehensive model that predicts mRNA degradation rates. Using multiple regression analysis, we find that 43% of variation in mRNA degradation rates determined using RATE-seq can be explained by considering multiple inherent properties of transcripts in a single model. By including association with specific RNA binding proteins and the function of the encoded product, ∼50% of the genome-wide variation in mRNA degradation rates can be explained. Interestingly, we find that methods for measuring RNA degradation that use transcriptional inhibition tend to explain far less variation than less disruptive methods. This may reflect that fact that metabolic labeling methods, which minimally perturb the cell, yield more physiologically relevant degradation rates than transcriptional inhibition, which results in rapid cell death.

In our analysis of mRNA degradation rates measured using RATE-seq, coding sequence length is the strongest single predictor of mRNA degradation rates: in general, the longer a transcript the more rapidly it is degraded. Other genome-wide investigations have shown a positive relationship between the length of the mature mRNA and its rate of degradation (46, 19, 47, 48). Studies of individual transcripts have shown that increasing transcript length by addition of specific sequences containing “instability elements” enhances a transcript’s rate of degradation (49). It is possible that it is not the length of the transcript that affects its degradation, but the presence of additional regulatory elements, to which *trans* factors such as RNA binding proteins can bind. A second, though not mutually exclusive explanation for the observed relationship with mRNA length, relates to the abundance of a transcript. The highest transcript levels are achieved through rapid synthesis and slow degradation. Therefore, the most abundant transcripts are expected to have slow rates of degradation. Within the transcriptome, the most abundant transcripts include those encoding ribosomal proteins and histones, which also tend to have the shortest coding length. The observed effect of length on mRNA degradation rates is likely a combination of both regulatory elements and non-random relationships between the encoded function of the transcript and its length.

### Association with ribosomes increases mRNA stability

Our multiple regression analysis provided evidence that protein translation impacts mRNA degradation rates. Previous studies regarding the role of translation on mRNA degradation have shown differing results. In a detailed study of the *CYC1* transcript, all start codons were removed from its coding region thereby preventing ribosome binding and translation (50). Following global transcription inhibition with thiolutin, the translationally impaired transcript degraded with similar kinetics to the translated transcript. Similarly, when translation of the *MFA2* transcript was inhibited by introducing a strong secondary structure in the 5’ region of the transcript it was not found to alter the degradation kinetics following transcriptional inhibition using the GAL system (51). By contrast, using the same method to reduce translation of the *PGK1* transcript, results in an increased mRNA degradation rate (6). Studies based on individual transcripts may be limited due to both interaction with additional factors and issues of statistical power. Our analysis shows that genome-wide, increased ribosome density is correlated with decreased mRNA rates (Figure 1 and Figure S3) as recently reported (42).

To study the effect of ribosome density on the rate of mRNA degradation we mutated the start codon of *GAP1* from an ATG to GTG. Consistent with results from our genome-wide study, loss of the first start codon results in reduced mRNA levels and an increased rate of mRNA degradation. Using a functional assay, we found that mutation of the start codon does not result in a loss of protein function (Figure S5). The next potential start codon is 288 nucleotides (96 amino acids) downstream of the wildtype start codon and thus we postulate that its use may result in a protein product that retains much of the wildtype GAP1 activity. Within the first 96 amino acids are residues known to affect the extent of functionality and localization of the encoded permease (52). Therefore, in addition to a decrease in ribosome density, we cannot exclude the possibility that alteration in protein function and/or possibly its production contributes to the decreased stability of the transcript.

### Synonymous coding mutations affect mRNA stability

Regression analysis suggested genome-wide relationships between codon usage and mRNA degradation rates. We find a negative correlation between codon adaptation index and mRNA degradation rate. Bias towards more frequent codons in a transcript may increase the rate of elongation during protein translation (53). Therefore, this negative correlation suggests that faster elongation by ribosomes may result in decreased rates of degradation. Consistent with this possibility we find that the presence of multiple sequential proline codons, which greatly slow the elongation rate, results in faster mRNA degradation rates.

We also find genome-wide evidence that the GC3 content is negatively correlated with rates of mRNA degradation. These results are consistent with an earlier study that found a positive correlation between synonymous A|T dinucleotides spanning codon boundaries and mRNA degradation rates (54). To validate this result, we experimentally tested the effect of GC3 content on mRNA degradation using GFP-encoding transcripts that differ in GC3 content. Consistent with our genome-wide analysis we find that decreasing the GC3 content results in increased mRNA degradation rates and lowered steady-state abundances. Recently, increased GC3 content has been suggested to decrease mRNA degradation rates in human lymphoblastoid cells and possibly explain some variation in mRNA degradation rates between different individuals (19). Thus, the relationship between GC3 content and mRNA degradation rates may be widely conserved in eukaryotes.

### Conclusion

Our study shows that genome-wide variation in mRNA degradation rates is best explained by a combination of different transcript features as suggested more than two decades ago (49). The fact that many properties of transcripts are correlated makes it difficult to identify causative relationships; however, through careful experimentation it is possible to confirm genome-wide principles as we have shown in this study. Many of the factors that affect genome-wide patterns of mRNA degradation rates are related to protein production highlighting the close relationship between mRNA degradation and translation, which warrants further investigation.

## ACKNOWLEDGEMENTS

We thank Joshua Plotkin and Grzegorz Kudla for providing the degenerate GFP constructs for our studies. We thank members of the Gresham Lab for helpful discussions and suggestions.

### AUTHOR CONTRIBUTIONS

B.N. performed all computational analyses and biological validation of GC content. V.E. performed biological validation of ribosome density. B.N. and D.G. conceived of the project and wrote the paper.

## SUPPLEMENTAL FIGURES

**Figure S1** Multiple predictors best explain variation in mRNA degradation. We show the adjusted R^2^ for individual predictors in each model. Predictors are as follows: l=Coding length, 2=3’UTR length, 3=5’UTR length, 4=5’UTR GC content, 5=3’UTR GC content, 6=mRNA abundance, 7=Protein per cell, 8=Ribosome density, 9=Transcription rate, 10=GC1, 11=GC2, 12=GC3, 13=Protein half-life, 14=deltaG, 15=CAI, 16=Protein per mRNA, 17=GO, 18=RNABP, 19=model with continuous variables, and 20=model with continuous variables +GO +RNABP.

**Figure S2** Predictors of mRNA degradation variation are highly correlated. We show that a number of predictors are highly correlated.

**Figure S3** Degradation rate decreases with increased ribosome density. The ribosome density of a transcript is inversely correlated with degradation rate, showing that the more ribosomes present, the more stable the transcript.

**Figure S4** GFP1 and GFP2 are similar in their degradation kinetics.

**Figure S5** Effect of mutated start codon on GFP protein. We grew yeast in minimal media containing D-histidine and D-serine. D-his is toxic to cells and leads to cell growth inhibition and death. In the absence of a functional GAP1 protein, cells do not have the permease and do not take up D-his. We show five strains: GAPl-WT=wild type strain, GAPl-KO=strain with complete knockout of coding sequence of GAP1, GAPl-NF=Derived strain with non-functional GAP1, GAPl-ATG=plasmid GAP1 with normal start codon, and GAPl-GTG=plasmid GAP1 with GTG in place of start codon.

**Figure S6** Diagnostics of the model fit.

## SUPPLEMENTAL TABLES

**Table S1** Matrix with all parameters included in regression analysis.

**Table S2** Matrix with p-value, adjusted R^2^, and correlation for each predictor and reported mRNA degradation rate.

**Table S3** Data for qPCR experiments.

